# Human variation impacting *MCOLN2* restricts *Salmonella* Typhi replication by magnesium deprivation

**DOI:** 10.1101/2022.05.08.491078

**Authors:** Kyle D. Gibbs, Liuyang Wang, Caroline E. Anderson, Jeffrey S. Bourgeois, Yanlu Cao, Margaret R. Gaggioli, Rosa Puertollano, Dennis C. Ko

## Abstract

Human genetic diversity can reveal critical factors in host-pathogen interactions. This is especially useful for human-restricted pathogens like *Salmonella enterica* serovar Typhi (*S*. Typhi), the cause of Typhoid fever. One key dynamic during infection is competition for nutrients: host cells attempt to restrict intracellular replication by depriving bacteria of key nutrients or delivering toxic metabolites in a process called nutritional immunity. Here, a cellular genome-wide association study of intracellular replication by *S*. Typhi in nearly a thousand cell lines from around the world—and extensive follow-up using intracellular *S*. Typhi transcriptomics and manipulation of magnesium concentrations—demonstrates that the divalent cation channel mucolipin-2 (MCOLN2) restricts *S*. Typhi intracellular replication through magnesium deprivation. Our results reveal natural diversity in Mg^2+^ limitation as a key component of nutritional immunity against *S*. Typhi.

**One-Sentence Summary:** Human immune cells genetically vary in their ability to use magnesium deprivation to restrict growth of the typhoid fever bacterium.

## Main Text

Genome-wide association studies (GWAS) are a powerful method to identify common variants associated with risk, resistance, or other quantitative measures of infectious disease. However, connecting GWAS variants from whole-organism studies to disease pathogenesis insight can prove challenging—especially when variants are near genes not clearly related to the disease under investigation. In this regard, GWAS of cellular traits, such as our platform Hi-HoST (<u>h</u>igh-throughput <u>h</u>uman *in vitr<u>o</u>* <u>s</u>usceptibility <u>t</u>esting) (*1, 2*), provide control of environmental and pathogen variation as well as facilitate subsequent mechanistic studies. Here, we used this approach to study susceptibility to the human-restricted enteric pathogen *Salmonella enterica* ser. Typhi (*S*. Typhi), which relies on a permissive niche inside immune cells to cause the life-threatening syndrome of Typhoid fever (*3*). We discovered that the interferon-inducible (*4*) host cation channel, mucolipin-2 (MCOLN2 or TRPML2), is a critical contributor to nutritional immunity against *S*. Typhi.

We identified human genetic variants associated with *S*. Typhi intracellular replication, using Hi-HoST screening and family-based GWAS analysis (*5*) of 961 lymphoblastoid cell lines (LCLs; EBV-immortalized B cells) from eight populations (**Fig. 1A and Data S1**). LCLs are a powerful *in vitro* model, because they are karyotypically normal, and B cells are a natural site of *Salmonella* replication *in vivo* (*6*). Intracellular replication, or host cell permissivity, is a demonstrated proxy for *Salmonella* virulence in whole organisms (*7*). In LCLs, we defined permissivity as the ratio of bacterial burden at 24 hours to 3.5 hours, based on median green fluorescence intensity of live intracellular *S*. Typhi. This analysis revealed a single genome-wide significant locus (rs10873679, p=6 ×10^−9^) on chromosome 1 (**Fig. 1B**). A quantile-quantile plot demonstrated no overall inflation of the test statistic, with primarily rs10873679-linked SNPs deviating from the neutral distribution (**Fig. 1C**). The association signal covers two genes in the mucolipin family, *MCOLN2* and *MCOLN3* (**Fig. 1D**). Mucolipins are a family of three inward rectifying divalent cation channels that localize to endolysosomal membranes and regulate vesicular trafficking (*8*).

**Figure 1.**
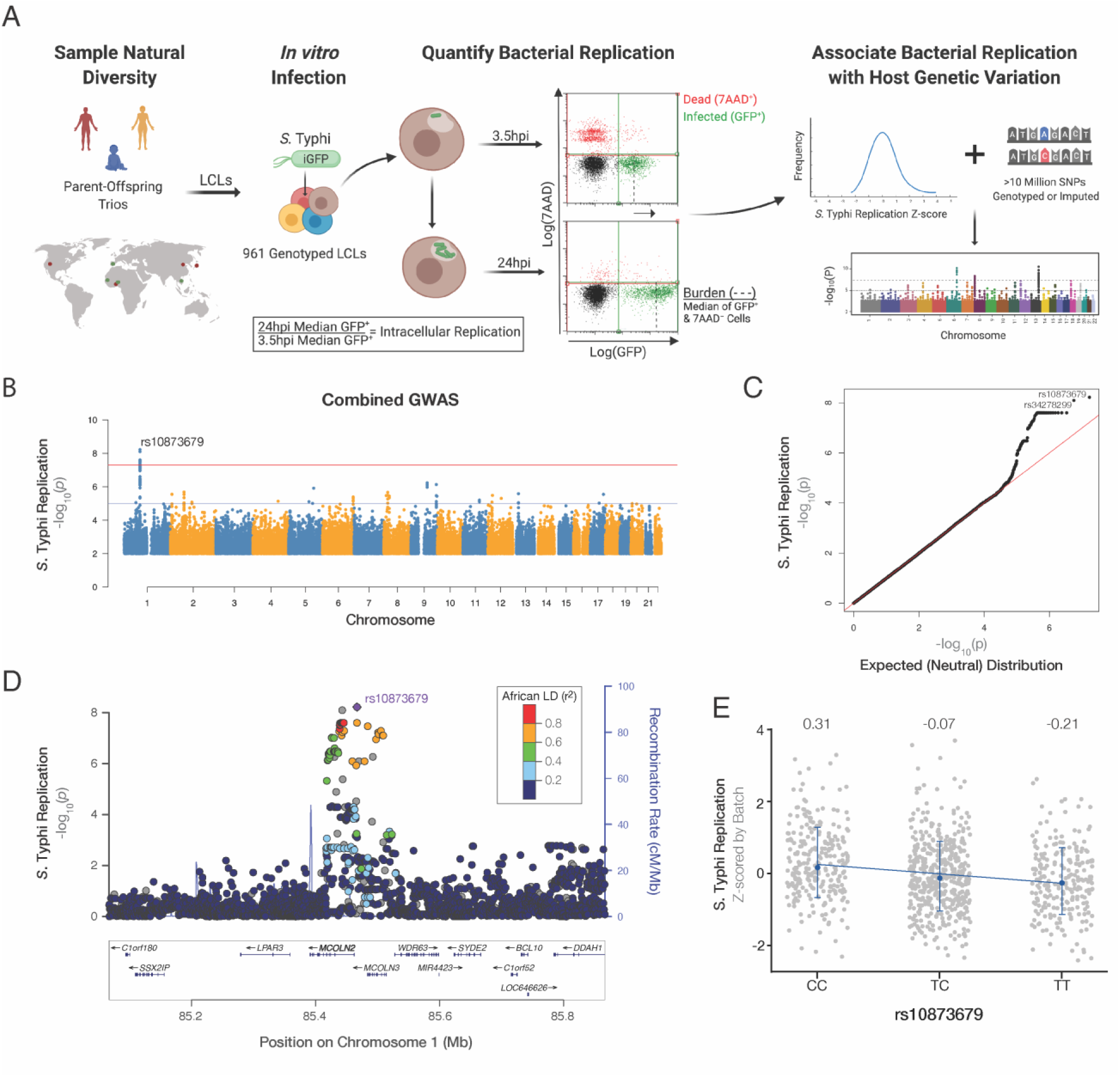
Cellular GWAS associates the rs10873679 locus with *S*. Typhi intracellular replication. (**A**) Hi-HoST cellular GWAS workflow. (**B**) Manhattan plot of cellular GWAS. P values were calculated using QFAM-parents on the z-scored replication ratios (orange line is p<5 ×10^−8^). The lead SNP on chromosome 1 is rs10873679 (p=6 ×10^−9^). (**C**) GWAS of *S*. Typhi intracellular replication has p values lower than expected from a neutral, χ^2^, distribution (red line). (**D**) A local Manhattan plot of the *S*. Typhi replication-associated locus on chromosome 1 with dots color coded by African linkage disequilibrium (LD; r^2^) from 1000 genomes Nov 2014 population build. (**E**) The rs10873679 C-allele is associated with increased *S*. Typhi replication. Means for each genotype are indicated above scatterplots. Bars are ±SD. Regression slope (β=-0.26±0.04) is significantly less than zero (p=1.7×10^−9^).

The minor C-allele of rs10873679 is associated with more intracellular replication (**Fig. 1E; Fig. S1**). To link this to cellular physiology, we examined expression of *MCOLN2* and *MCOLN3* in RNA-seq of 1000 Genomes LCLs (*9*). The C-allele associated with less *MCOLN2* expression (**Fig. 2A**; p<2×10^−16^), while rs10873679 was not associated with a significant difference in *MCOLN3* expression (**Fig. S2**). In confirmation, the C-allele also associated with reduced MCOLN2 protein abundance (*10*) (**Fig. 2B**). The rs10873679 C-allele’s association with both more *S*. Typhi replication and less *MCOLN2* expression suggested that MCOLN2 restricts *S*. Typhi intracellular replication.

**Figure 2.**
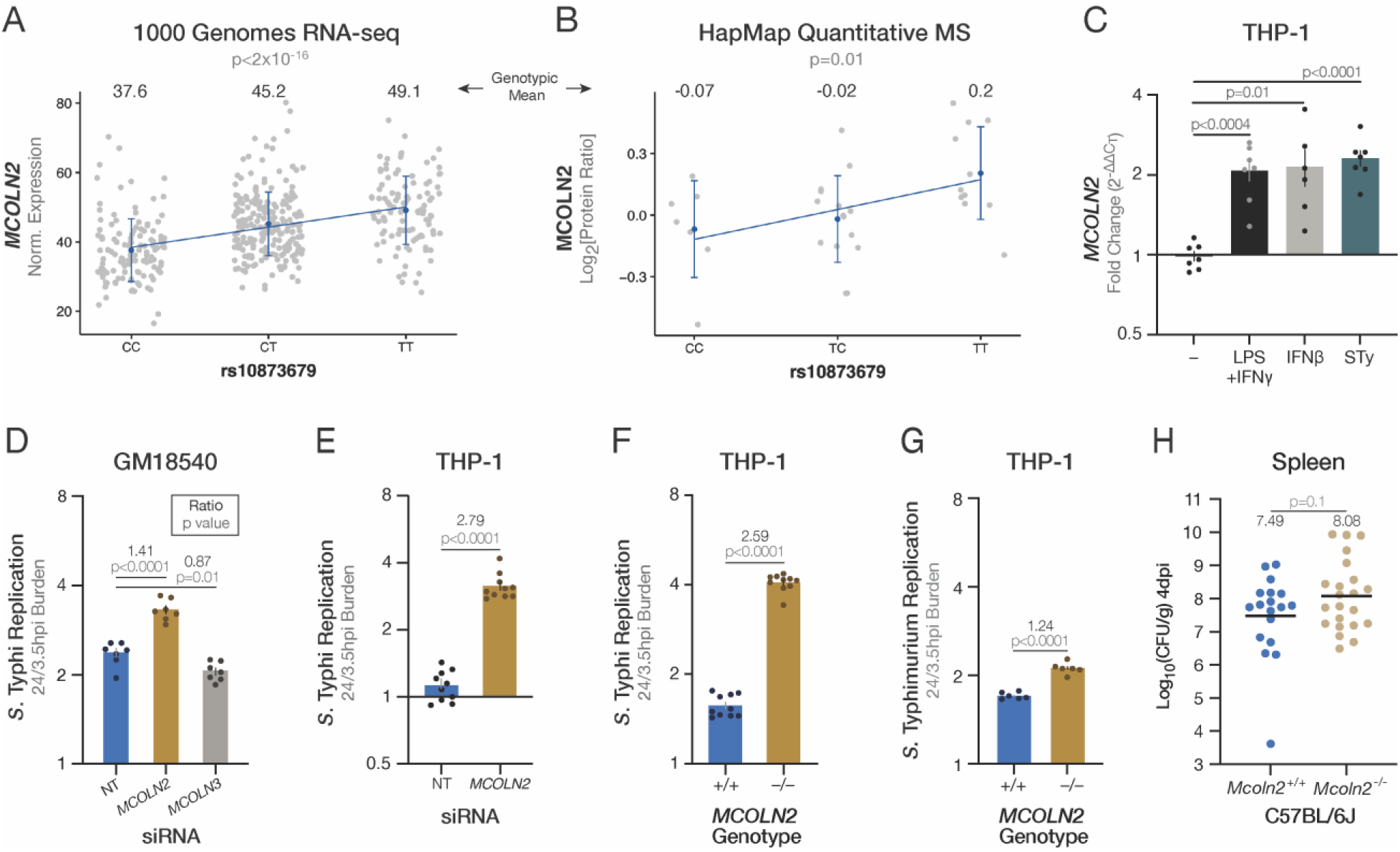
Increased mucolipin-2 expression restricts *S*. Typhi replication in human immune cell lines. (**A**) The rs10873679 C-allele associates with less *MCOLN2* mRNA expression in 1000 Genomes Project LCLs (*9*). Linear regression of 448 LCLs is significant (β = 5.8±0.6; p <2 ×10^−16^; r^2^ = 0.166). (**B**) The rs10873679 C-allele associates with less MCOLN2 protein expression in 33 LCLs measured with mass spectrometry (*10*). Linear regression is significant (β = 0.15 ±0.05, p=0.01; r^2^=0.017). Bars in A-B are mean ±SD. (**C**) Both interferon treatment and *S*. Typhi (STy) infection (MOI 10, 24 hrs) stimulate *MCOLN2* expression in THP-1s. Expression measured by RT-qPCR and quantified by ΔΔC_T_ (ΔC_T_ stimulated - ΔC_T_ untreated). Seven replicates from three experiments. P values are from Dunnet’s T3 comparison after Welch’s ANOVA (p<0.0001). (**D**) *MCOLN2* knockdown increases *S*. Typhi replication, while *MCOLN3* knockdown modestly decreases *S*. Typhi replication in comparison to non-targeting (NT) siRNA. Seven replicates from three experiments. Knockdown qPCR: 0.33-fold (±0.14) of NT *MCOLN2* expression and 0.53-fold (±0.04) of NT *MCOLN3* expression. P values are from Dunnett post-hoc comparison to NT following a one-way ANOVA (main effect p<0.0001). (**E**) *MCOLN2* knockdown (0.09-fold (±0.02) of NT) increases *S*. Typhi replication in THP-1s. Ten replicates from two experiments. (**F**) CRISPR-Cas9 knockout of *MCOLN2* increases *S*. Typhi replication in THP-1s. Ten replicates from two experiments. (**G**) *S*. Typhimurium has a minor growth advantage in *MCOLN2* knockout THP-1s. Six replicates from two experiments. In D-G, ratios are mean in siRNA-treated/NT or knockout/wild-type cells. In C-G, bars are mean ±SEM and all statistics are calculated with log_2_-transformed data. (**H**) *Mcoln2* knockout does not significantly increase burden in C57BL/6J mice spleens four days post IP infection with 1,000 CFUs of late-log *S*. Typhimurium (14028s) tagged with p67GFP3.1. 18 wild types and 22 knockouts from six experiments. Lines are geometric means and log_10_(geo. mean) is written above each genotype. P value calculated with log_10_-transformed data. Without the low outlier (Log_10_[CFU]=3.6; identified at Grubbs’ α=0.01), the *Mcoln2*^+/+^ log_10_(geo. mean) is 7.72 and p=0.2. In E-H, p values are from Welch’s t test.

Strengthening this model, *MCOLN2* is upregulated in human macrophages after treatment with M1 polarizing LPS & IFN-γ (*11*), which indicates that *MCOLN2* is part of the host response. Similarly, we observed *MCOLN2* induction after *S*. Typhi infection (**Fig. 2C**). If MCOLN2 is a restriction factor, we expected ablating *MCOLN2* expression would increase intracellular *Salmonella* replication. Knocking down *MCOLN2*, but not *MCOLN3*, increased *S*. Typhi intracellular replication (**Fig. 2D**), without affecting bacterial invasion or pyroptosis (**Fig. S3**). This phenotype generalized to other human immune cells, as knocking down *MCOLN2* in THP-1 monocytes by RNAi (**Fig. 2E**) or knocking out the gene using CRISPR/Cas (**Fig. 2F**) resulted in an even greater increase in *S*. Typhi replication than in LCLs.

rs10873679 was also associated with intracellular replication of *S*. Typhimurium (p=8.1×10^−7^; **Fig. S4**), a serovar used to model enteric fever in mice as *S*. Typhi is human-restricted; however, the impact of reducing *MCOLN2* expression is much smaller with *S*. Typhimurium (**Fig. 2G**). This demonstrates that while MCOLN2 is a key restriction factor for *S*. Typhi (knockout results in ∼150% more replication), it is an accessory factor for controlling *S*. Typhimurium (knockout results in ∼20% more replication). We confirmed lack of a large effect with *S*. Typhimurium by infecting susceptible C57BL/6J mice with *Mcoln2* knocked out (*12*) via intraperitoneal injection—which avoids restriction by stomach acid or variance introduced by gut microbiota—and quantified *S*. Typhimurium burden in the spleen four days post infection (**Fig. 2H**). This revealed no significant difference in *S*. Typhimurium burden between *Mcoln2* genotypes, despite a modest trend of higher burden in *Mcoln2*^*-/-*^ mice, which is not surprising given the small *in vitro* phenotype. This serovar difference could be explained by bacterial difference—only *S*. Typhi has the capacity to take advantage of a changed niche after MCOLN2’s removal—or a differential host response, in which the more immunogenic *S*. Typhimurium induces additional restriction factors that prevent it from fully exploiting *MCOLN2* knockout. Regardless, our data demonstrate that MCOLN2 is a strong restriction factor for the human-specific serovar *S*. Typhi, which underscores the value of cellular GWAS for identifying human-specific host-pathogens interactions.

To determine how MCOLN2 reduces *S*. Typhi replication, we used the intracellular bacteria as reporters of their own environment. We conducted transcriptomics at 16 hours post infection (hpi), near maximum divergence of replication inside wild-type versus *MCOLN2*^*-/-*^ THP-1s and prior to restriction in wild-type THP-1s (**Fig. 3A & B**). While >2,600 bacterial genes were detected, and expression of one quarter of the bacterial transcriptome significantly changed between late-log inoculum and 16 hpi, differences between bacteria within wild-type and *MCOLN2*^*-/-*^ cells were more modest with expression of no individual bacterial gene passing significance threshold after correction for multiple testing (**Data S2**). Therefore, we used gene set enrichment analysis (GSEA) to identify *S*. Typhi processes that were upregulated in *MCOLN2*-containing wild-type THP-1s. We generated a list of 15 genes sets of physiological processes associated with virulence or divalent cation transport (**Fig. 3C and Table S1**). Only genes regulated by the PhoP/Q two-component system (*13*) were significantly enriched (NES = - 1.81 with FDR q=0.004) in bacteria living inside wild-type THP-1s compared to *MCOLN2*^-/-^ THP-1s (**Fig. 3D**). While *S*. Typhi within *MCOLN2*^*-/-*^ cells upregulate PhoP/Q targets (10.7-fold more expression than late-log), induction is greater in bacteria inside wild-type cells (13.3-fold).

**Figure 3.**
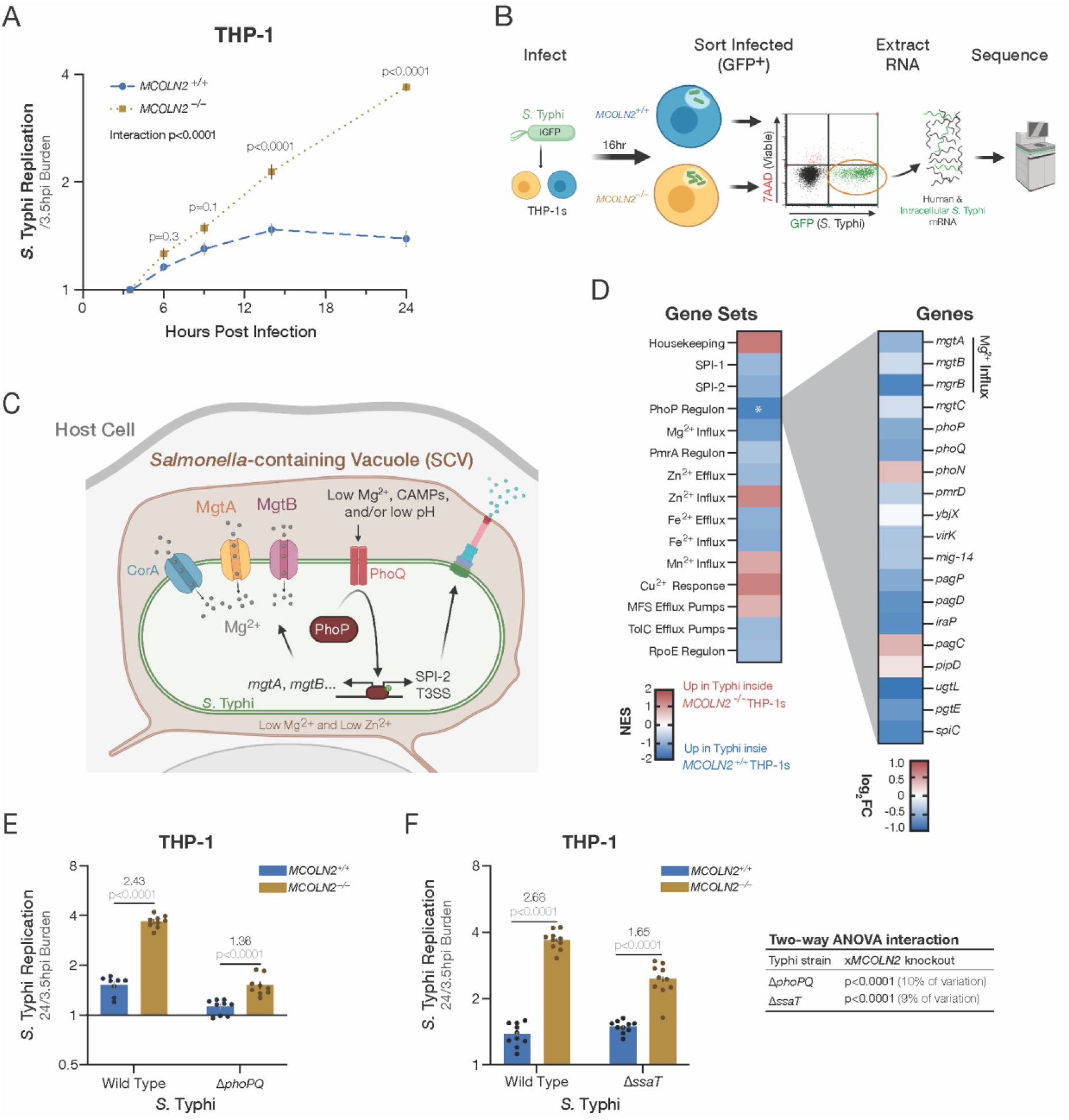
Intracellular *S*. Typhi’s replication inside *MCOLN2*^-/-^ THP-1s depends on PhoP/Q. (**A**) *MCOLN2* knockout leads to faster *S*. Typhi replication inside THP-1s. Ten replicates from three experiments, except 6 hpi is six replicates from two experiments. *MCOLN2* genotype, hpi, and their interaction are all significant sources of variation (p<0.0001) in a repeated measures ANOVA. Time point p values are from Šídák’s post-hoc comparison of *MCOLN2*^+/+^ and *MCOLN2*^−/–^. (**B**) Workflow used to sequence mRNA from intracellular *S*. Typhi 16 hpi in wild-type and knock out THP-1s. (**C**) Diagram of *S*. Typhi’s PhoPQ-induced Mg^2+^ importers. (**D**) RNA-seq of intracellular *S*. Typhi indicates PhoP targets are upregulated more when MCOLN2 is present. Left: Normalized enrichment score (NES) from GSEA of virulence-or cation-associated *S*. Typhi gene sets. Significant gene set (FDR q<0.05) indicated by asterisk. Right: The log_2_(KO/WT expression) of PhoPQ regulon genes are plotted. 16 of 19 genes have a negative fold-change (FC), indicating higher expression in WT. (**E**) *PhoPQ* is required for most of the increase in intracellular replication observed with *MCOLN2*^*-/-*^ THP-1s. Nine replicates from two experiments. (**F**) *S*. Typhi Δ*ssaT* has no effect on intracellular replication in WT THP-1s and partially accounts for the requirement of *phoPQ* to achieve maximal replication in *MCOLN2*^*-/-*^ THP-1s. Ten replicates from three experiments. P values in E & F are from Šídák’s comparison of *MCOLN2*^+/+^ to *MCOLN2*^−/–^ following two-way ANOVAs finding significant main effects and interaction (all p<0.0001). Statistics in A, E, & F use log_2_-transformed replication ratios. Bars in A, E, & F are mean ±SEM.

To determine if PhoP/Q signaling contributes to replication in *MCOLN2* knockout cells, we infected THP-1s with the Ty800 Δ*phoPQ* strain (*14*), which revealed that most (∼75%) of the increased replication inside *MCOLN2*^*-/-*^ requires intact PhoPQ signaling (**Fig 3E**). Chief among PhoP/Q targets is the SPI-2 T3SS that injects effectors to maintain *Salmonella*’s intracellular niche. Removing an essential component of the SPI-2 T3SS basal body (*ssaT*) to prevent any secretion caused no change in *S*. Typhi replication within wild-type THP-1s (compare blue bars in **Fig. 3F**). This contrasts to *S*. Typhimurium (*15, 16*) but is consistent with past *S*. Typhi literature (*17*). In contrast, replication is reduced in *MCOLN2*^*-/-*^ cells, suggesting that roughly half of the PhoP/Q-dependent increase in *S*. Typhi replication depends on SPI-2 effectors (**Fig. 3F**). This indicates the SPI-2 independence of *S*. Typhi replication in THP-1 monocytes is actually an MCOLN2-dependent host response that suppresses the fitness advantage provided by *S*. Typhi’s SPI-2 effectors.

Our results demonstrate that *S*. Typhi replicating inside *MCOLN2*^-/-^ monocytes upregulate PhoP targets, which significantly boosts replication. However, in wild-type cells, the even greater induction of PhoP targets is not sufficient to increase replication, so we speculated the PhoP upregulation was a symptom of a restrictive condition enhanced by MCOLN2. Three potentially restricting conditions in the SCV lead to more PhoP activity: PhoP/Q is repressed by high Mg^2+^(*18*) and activated by cationic antimicrobial peptides (CAMPs) (*19*) or acidification (*20, 21*). Since MCOLN2 is a divalent cation channel, PhoP/Q was most likely responding to reduced Mg^2+^ concentrations, which along with Zn^2+^, are limited in SCVs (*22, 23*). Indeed, the PhoP-activated Mg^2+^ importers *mgtA* and *mgtB* were both upregulated more in bacteria inside wild-type THP-1s (**Fig. 3D; Fig. S5**). Therefore, the transcriptomics and *phoPQ* mutant infection data suggested a simple hypothesis: MCOLN2 deprives *S*. Typhi of Mg^2+^. To test this, we repleted Mg^2+^ two hours after infecting and measured bacterial replication (**Fig. 4A**). Mg^2+^ supplementation disproportionately benefited bacterial replication inside wild-type THP-1s (1.6-fold in wild-type vs. 1.2-fold in knockout THP-1s; interaction p=0.002). Similar results were also observed with *S*. Typhimurium (**Fig. S6**). While our transcriptomics could also support a role for Zn^2+^, zinc repletion did not have interactions with *MCOLN2* genotype (**Fig. 4B**; interaction p=0.3). Together, these data demonstrate that intracellular replication is held back by magnesium starvation and not zinc toxicity.

**Figure 4.**
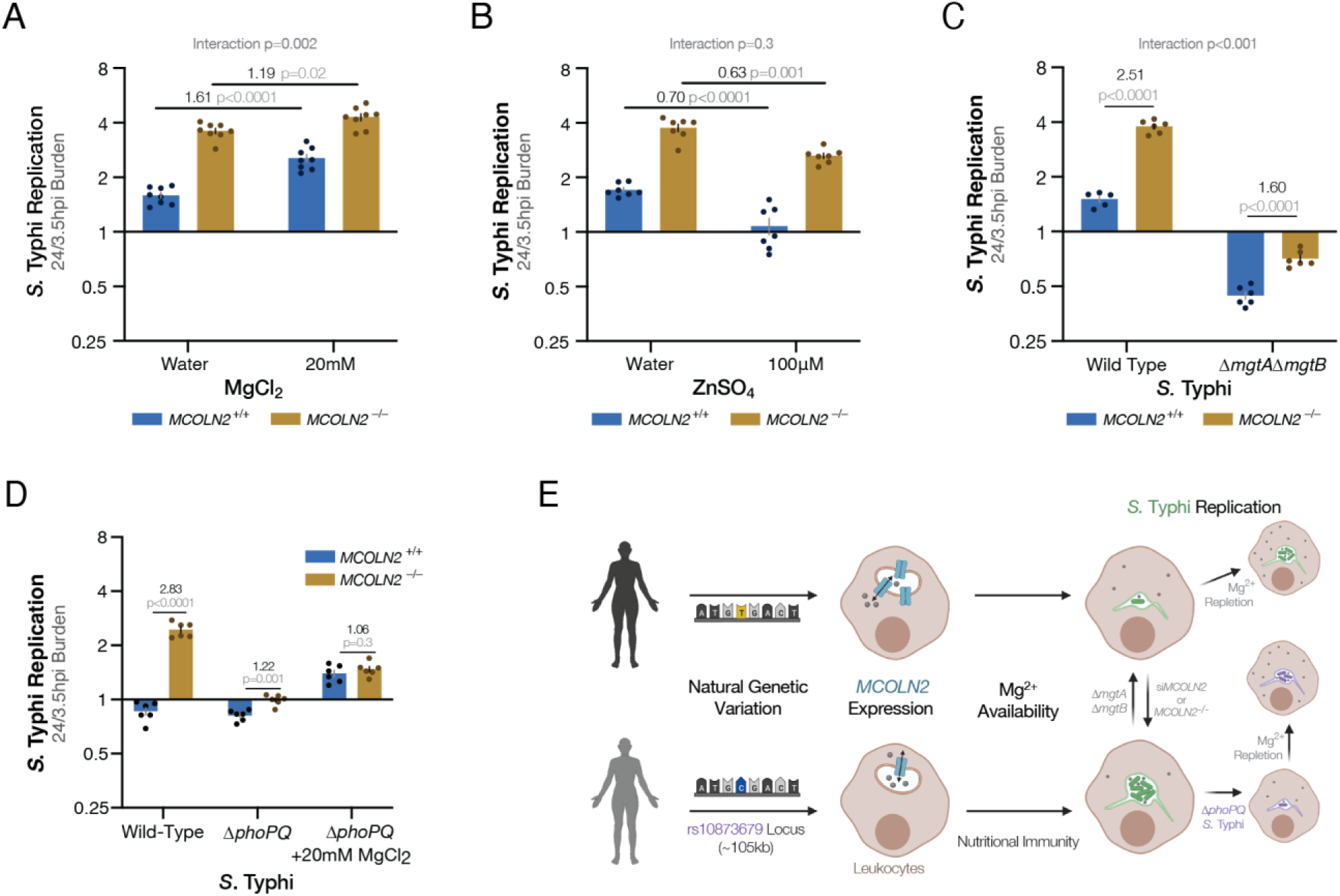
MCOLN2 reduces *Salmonella* replication by reducing magnesium availability. (**A**) Mg^2+^ supplementation partially rescues *S*. Typhi replication in *MCOLN2*^-/-^ THP-1s. Eight replicates from two experiments. Mg^2+^ supplementation, *MCOLN2* genotype are both significant (p<0.0001) in two-way ANOVA. (**B**) Zn^2+^ supplementation reduces *S*. Typhi replication in THP-1 monocytes independently from *MCOLN2* genotype. Seven replicates from two experiments. In a two-way ANOVA, Zn^2+^ supplementation and *MCOLN2* genotype are significant sources of variation (p<0.0001), but their interaction is not (p=0.3). (**C**) PhoPQ-induced magnesium importers MgtA and MgtB are required for half the *S*. Typhi replication benefit in *MCOLN2*^-/-^ THP-1s. Six replicates from two experiments. In a two-way ANOVA, *MCOLN2* genotype, *mgtA*/*mgtB* deletion, and their interaction are all significant sources of variation (p<0.0001). In A-C, p values comparing two means are from post-hoc Šidák’s multiple comparison tests. (**D**) *MCOLN2* knockout does not benefit Δ*phoPQ S*. Typhi replication inside THP-1s when Mg^2+^ is repleted. Six replicates from two experiments. In a three-way ANOVA, MgCl_2_ treatment, *phoPQ* deletion, *MCOLN2* genotype, and all two-way interactions are significant (p<0.0001). P values comparing two means are from post-hoc Welch’s t tests. (**E**) Model of natural genetic variation altering intracellular *S*. Typhi replication.

The importance of Mg^2+^ acquisition for *Salmonella* replication is underscored by its trio of Mg^2+^ uptake proteins: one constitutive, CorA, and two inducible, MgtA & MgtB. If knocking out *MCOLN2* increases Mg^2+^ availability, we theorized that these transporters would be necessary to uptake that extra Mg^2+^ and therefore essential for the enhanced replication inside *MCOLN2*^-/-^ host cells. To test this, we generated a double knockout (Δ*mgtA*Δ*mgtB*), which lacks the high affinity Mg^2+^ importers used in low-Mg^2+^ environments, like the ≤10μM concentration in the SCV (*22*). The double importer mutant is killed, instead of replicating, inside THP-1s (**Fig. 4C**). And in confirmation of our theory, knocking out *MCOLN2* provides less of an advantage to the double importer mutant (increasing replication 60% in ΔΔ vs. 150% in wild-type *S*. Typhi; interaction p<0.001). This corroborates the Mg^2+^ repletion and suggests that most of the enhanced replication in *MCOLN2*^-/-^ THP-1s depends on increasing Mg^2+^ availability. In the simplest version of our model, removing MCOLN2 increases Mg^2+^ availability to *S*. Typhi, which relieves a nutrient limitation and directly increases bacterial replication. However, ∼1/3 of the increased bacterial replication inside *MCOLN2* knockout cells is not explained by manipulating Mg^2+^ availability or uptake. We theorized that this putatively Mg^2+^-independent replication boost in *MCOLN2*^-/-^ cells could still be PhoP regulated, as we had already identified other PhoP-targets, namely SPI-2 T3SS, which further benefit *S*. Typhi replication when *MCOLN2* is knocked out. To test this, we repleted Mg^2+^ after infecting THP-1s with *S*. Typhi Δ*phoPQ* (**Fig. 4D**). Mg^2+^ increased replication of Δ*phoPQ* bacteria, as it partially overcomes the inability to fully upregulate *mgtA* and *mgtB*. Further, the combined Mg^2+^ repletion and *phoPQ* deletion removed any discernable difference in *S*. Typhi replication between *MCOLN2* genotypes. Thus, enhanced bacterial replication in the absence of *MCOLN2* depends on both Mg-independent effects of PhoPQ and PhoPQ-independent effects of Mg^2+^ availability (**Fig. 4E**).

In this report, we directly connect the divalent cation channel MCOLN2 with variable immune cell permissivity to *S*. Typhi. For *S*. Typhimurium, intracellular replication regulates outcomes in mouse models of enteric fever (*7, 24*), and therefore, *S*. Typhi replication likely also correlates with disease outcome in humans. Unfortunately, there is no published GWAS of typhoid severity or clinical outcome, and only one study on Typhoid fever onset, which identified an association between the MHC region and susceptibility (*25*). Thus, determining the clinical significance of rs10873679 in humans awaits well-powered studies for this disease phenotype. Our findings also underscore that despite great insights gleaned from mouse models of *S*. Typhimurium infection, studies of genetic diversity using human-specific pathogens in human cells provide unique insight. We showed that *MCOLN2* ablation reduced the low-Mg^2+^ stress faced by intracellular *S*. Typhi based on lowered expression of PhoP targets, including key Mg^2+^ transporters, and reduced benefit of Mg^2+^ repletion. This suggests that MCOLN2 exerts a similar restriction pressure on *S*. Typhi inside human monocytes as the divalent cation transporter Slc11a1 (Nramp1) does on *S*. Typhimurium inside murine macrophages (*26*). While to our knowledge, no paper has shown SLC11A1 restricting *S*. Typhi growth in human cells, we propose that MCOLN2 is preforming a functionally analogous role by also reducing Mg^2+^ availability to *Salmonella* in a different host-serovar pair.

Identifying MCOLN2 as another host factor that reduces *Salmonella* replication by lowering Mg^2+^ availability highlights the key role played by Mg^2+^ in nutritional immunity. This builds on a line of work identifying the sophisticated regulatory network in *S*. Typhimurium that allows it to respond to the low-Mg^2+^ environment of the SCV (*27, 28*). Notably, these investigations into *Salmonella* response to low Mg^2+^ have been conducted with non-typhoidal *S*. Typhimurium. While much of this regulatory system is likely preserved in *S*. Typhi, the much greater sensitivity of *S*. Typhi to MCOLN2 ablation suggests that some component of this low Mg^2+^ response is not conserved between the serovars. Future studies investigating this difference could reveal key serovar-specific virulence strategies.

Our finding that MCOLN2 restricts *S*. Typhi also explains why it is an ISG, despite previous findings that it increases macrophage susceptibility to endocytosed viruses including influenza A virus (*Orthomyxoviridae*) and yellow fever virus (*Flaviviridae*) (*29*). This identifies the *MCOLN2* locus as a possible site of balancing selection between different infectious disease pressures—viruses that use the endocytic pathway for entry might select for people with less *MCOLN2* expression, while Salmonellae infections might select for people with more *MCOLN2* expression. This balancing selection could explain the wide distribution of both rs10873679 alleles in populations around the world, and ultimately, highlights the persistent and complex power of infectious disease as an evolutionary pressure shaping human evolution.

## Materials and Methods

### Cell Culture

Lymphoblastoid cell lines (LCLs; EBV-immortalized B cells) were from the Coriell Institute. *MCOLN2*^-/-^ and matched wild-type THP-1 cell pools were generated by Synthego using guide 5’- ttttggtttaagtaaccagc-3’ (PAM is TGG) to target the start of *MCOLN2* exon 3. THP-1 knock out pools were confirmed to maintain ≥85% frameshift indels by Sanger sequencing that was analyzed with the inference of CRISPR editing (ICE) webtool v2.0 (https://ice.synthego.com) from Synthego(*30*). THP-1s and LCLs were maintained at 37°C in a 5% CO_2_ atmosphere and were grown in RPMI 1640 media (Gibco #21870) supplemented with 10% heat-inactivated fetal bovine serum (HI-FBS, Gibco #10082), 2 mM L-glutamine (Gibco #25030081), &100 U/ml Penicillin-Streptomycin (Gibco #15140122). Infection assays were carried out in the same media but without Pen-Strep and phenol red. Cells were verified as mycoplasma free by the Universal Mycoplasma Detection Kit (ATCC #30-1012K).

### Bacteria Strains

*Salmonella enterica* serovars Typhi strain Ty2, Typhimurium strain 14028s, and derived mutants, see Table S2, were grown at 37°C and 250 rpm in high-salt Miller Luria-Bertani (LB) broth (VWR #90003). To quantify intracellular burden during gentamicin protection assays, Salmonellae were tagged with inducible GFP using p67GFP3.1 (*31*), which carries GFP under an IPTG-inducible promoter and is maintained with 100 mg/mL ampicillin.

*Salmonella* gene deletion strains, listed in Table S2, were generated by lambda-red recombineering (*32*) from Ty2 or 14028s using Kan^R^ cassettes generated from pKD4 with the primers in Table S3. Gene deletions were confirmed by PCR using indicated primers in Table S3.

### Infection (Gentamicin-protection) Assays

*Salmonella* infection of LCL and THP-1 cells was done as previously described (*33*). In brief, overnight stationary cultures in Miller LB were sub-cultured 1:33 and grown for 160 minutes at 37°C and 250 rpm to reach SPI-1 inducing late-log phase (an OD_600_ of 1.7–2.0 for *S*. Typhimurium and 0.8–1.1 for *S*. Typhi). 1×10^5^ LCL or THP-1 cells were plated at 1×10^6^ cells/mL in complete RPMI one hour before infection in 96-well non-TC plates. LCLs were infected at multiplicity of infection (MOI) 30 and THP-1s at MOI 10. To kill the extracellular bacteria, gentamicin was added one hour post infection (hpi) at 50 mg/mL and then diluted to 15 mg/mL at 2 hpi. In ion repletion experiments, 5μL of filter-sterilized MgCl_2_ or ZnSO_4_ in DI water, or water only control, was added to 200μL in 96-well plates immediately following gentamicin dilution at 2 hpi. To induce GFP in p67GFP3.1, 1.4 mM IPTG was added 75 min prior to the desired time point.

Invasion, pyroptosis, and initial burden were measured with a Guava EasyCyte Plus high-throughput flow cytometer (Millipore) at 3.5 hpi. Pyroptosis was quantified as the percent staining with 1 μg/mL 7AAD (7-aminoactinomycin D; Enzo Life Sciences). Invasion was quantified as the percent GFP^+^ & 7AAD^-^. Burden was quantified as median fluorescent intensity (MFI) of living (7AAD^−^) and infected (GFP^+^) cells. Intracellular replication (permissivity) was quantified by re-measuring burden at 24 hpi and taking the ratio of 24 hpi burden over initial 3.5 hpi burden.

### Cellular GWAS

Hi-HoST screening of 961 LCLs from parent-offspring trios for *S*. Typhi intracellular replication occurred in two large sets. In one, *S*. Typhi intracellular replication was one of 79 host-pathogen phenotypes measured as part of the Hi-HoST Phenome Project (H2P2) (*2*). H2P2 measured replication in 527 LCLs from four population in the 1000 Genomes Project (*34*): ESN (Esan in Nigeria), GWD (Gambians in Western Divisions in The Gambia), IBS (Iberian Population in Spain), and KHV (Kinh in Ho Chi Minh City, Vietnam). In this dataset, we determined that replication is a quantitative trait suitable for GWAS due to its interindividual variation (mean of 1.7-fold with standard deviation of 0.3), high experimental repeatability (∼75% variance is due to inter-individual variation in two-way ANOVA), and substantial heritability (h^2^=0.33 with p=0.002 in parent-offspring regression) (*2*). To these 527 LCLs, we added previously unpublished data on *S*. Typhi replication from 434 LCLs from four populations in the HapMap project: CEU (Utah residents with ancestry from northern and western Europe), YRI (Yoruba in Ibadan, Nigeria), CHB (Han Chinese in Beijing, China), and JPT (Japanese in Tokyo, Japan) (*35*). For all 961 LCLs, we used flow cytometry to quantify intracellular bacterial burden as the median fluorescent intensity (MFI) of GFP in infected host cells, which contain viable GFP-tagged *S*. Typhi (see above for details of this fluorescence-based gentamicin protection assay). From these MFI measurements, we calculated intracellular replication or permissivity as the ratio of 24 hpi to 3.5 hpi burden. Each LCL was measured on three sequential passages and the phenotype used for GWAS was calculated as the mean measurement of these three independent assays. Each batch of LCLs measured during Hi-HoST screening was z-score transformed to reduce inter-batch experimental variation: *Z* = (*x* − *μ*_*batch*_)/*σ*_*batch*_.

Genotypes were obtained from HapMap r28 and 1000 Genomes Project Phase 3 with imputation using 1000 Genomes Project Phase 3. Filters included minor allele frequency (MAF) < 0.05, SNP missingness of > 0.2 and sample genotype missingness of > 0.2, resulting in a total of 8,386,469 SNPs for subsequent analysis. Genome-wide association analysis was carried out using the QFAM-parents approach in PLINK v1.9 (*5, 36*) with adaptive permutations ranging from 1000 to a maximum of 10^9^. The QFAM approach uses linear regression to test for association while separately permuting between and within family components to control for family structure. The human genome reference assembly (GRCh37/hg19) was used for all analysis. Intracellular replication data for the 961 LCL samples can be found in Data S1 and GWAS summary statistics are available for download at the Duke Research Data Repository (https://doi.org/10.7924/r4x92bd76). QQ plots against neutral, χ^2^, distribution were plotted using quantile-quantile function in R. Local Manhattan plots were generated using LocusZoom (*37*) webtool (http://locuszoom.org/). Linear regression of *Salmonella* replication by rs10873679 genotype was performed and plotted in R using ggplot2 (*38*) & ggthemes (*39*) packages.

### Human Gene Expression Analyses

RNA-seq gene expression data of 448 LCLs from the 1000 Genomes Project (*40*) were obtained from the EBI website (https://www.ebi.ac.uk/gxa/experiments/E-GEUV-1/Downloads). The rs10873679 genotype data were downloaded from the 1000 genome project (*41*). Effects of rs1087369 genotype on *MCOLN2* and *MCOLN3* gene expression in both datasets were tested by linear regression on combined data as well as individual populations and individual sexes.

Protein abundance measured by isobaric tag-based quantitative mass-spectroscopy in 95 LCLs from HapMap project were obtained from Wu *et al (42)*. However, only 33 of the individuals had quantifiable MCOLN2. The effect of rs10873679 on MCOLN2 protein abundance was tested by linear regression in R.

### RNAi Experiments and Knockdown Confirmation

LCLs or THP-1s (2.5 ×10^5^ cells) were washed and re-suspended at 400,000 cells/mL in 500 μL of serum-free Accell siRNA delivery media (Horizon #B-005000) in TC-treated 24-well plates and treated for three days with 10 pg/μL of either Dharmacon Accell non-targeting #1 (NT1) (Horizon #D-001910-01) or an Accell SmartPool against human *MCOLN2* (Horizon #E-021616-00) or *MCOLN3* (Horizon #E-015371-00). Prior to infection, cells were washed and plated at 700,000 cells/mL in 100 μL complete RPMI media (without antibiotics) in 96-well non-TC plates. Infections were conducted as described above.

For each experiment, knockdown was confirmed by RT-qPCR. Briefly, RNA was extracted from one well not used for infection (∼5×10^5^ treated cells) for each siRNA condition using RNeasy kit (Qiagen #74106). Then cDNA was reverse transcribed from 500ng of RNA/condition using iScript kit (BioRad #1708891) and quantified by qPCR using iTaq Universal Probes Supermix (BioRad #1725134) and exon-spanning TaqMan FAM-MGB probes (ThermoFisher #4331182; *MCOLN2* is Hs00401920 & *MCOLN3* is Hs00962657) on a QuantStudio 3 thermocycler (ThermoFisher). All qPCR was run in technical triplicate. Mean comparative threshold cycle (C_T_) value for each transcript was adjusted for input variation by subtracting the mean 18s (*RNA18S5*; ThermoFisher Hs03928990) housekeeping control C_T_ from the target gene’s C_T_ to generate a ΔC_T_. The ΔΔC_T_ for each knockdown was calculated by subtracting target gene ΔC_T_ in siNT1-treated control cells from target gene ΔC_T_ in siTarget-treated cells. Knockdown fold change was then calculated as 2^-ΔΔCT^. Mean fold-change knockdown ±SEM was reported in figure legends.

### Inducing and Measuring *MCOLN2* Expression

To measure *MCOLN2* induction, 5×10^5^ THP-1s in 24-well non-TC treated plates were infected with *S*. Typhi Ty2 at MOI 10 following the above gentamicin-protection assay or stimulated 2 hpi with 500 U/mL (25ng/mL) recombinant human IFN-γ (PeproTech #300-02) and 100 pg/mL well-vortexed *S*. Typhimurium S-form LPS (Enzo #ALX-581-011) or 50 U/mL (5ng/mL) recombinant human IFN-β (PeproTech #300-02BC). At 24hpi, *MCOLN2* expression was measured by RT-qPCR following the same ΔΔC_T_ method used to measure knockdown.

### Fluorescent Activated Cell Sorting (FACS)

For cell sorting RNA-seq samples, 20 million THP-1 monocytes of each *MCOLN2* genotype were plated into 24-well non-TC plates (500,000 cells per 0.5 mL RPMI per well) and infected with *S*. Typhi at MOI10. The remaining late-log *S*. Typhi inoculum was washed with PBS and fixed with 100 μL of RNA*later* Solution (ThermoFisher #AM7020) for 10 min at room temperature and then frozen for later RNA extraction. Following 2 hours of IPTG induction, monocytes were spun down at 16 hpi and re-suspended at 10 million cells/mL in 2 mL of RPMI containing 15 μg/mL gentamicin and 1 μg/mL 7AAD. Two wells containing one million uninfected THP-1 monocytes of each genotype were washed with PBS and fixed in 1mL RNA*later* for use in the control.

Live monocytes were analyzed and sorted by the Duke Human Vaccine Institute (DHVI) flow cytometry shared resource using a FACSAria II (BD Biosciences) at 70 psi with at 70 μm nozzle. One million infected (GFP^+^) and living (7AAD^-^) cells were sorted into 1 mL of RNA*later* for immediate fixation and held at 4°C in a chilled collection tube rack. Doublets were excluded by FSC and SSC gating and a purity mask was applied to exclude droplets containing GFP^+^ and GFP^-^ events.

### *S*. Typhi infected THP-1 RNA Extraction

After sorting, RNA was immediately isolated from collected cells using mirVana miRNA Isolation Kit’s total RNA protocol (ThermoFisher #AM1560). Prior to extraction, samples in RNA*later* were diluted 3x with PBS, spun down at 5,000 xg, and aspirated to remove RNA*later*. After resuspending cells in the kit’s L/B buffer, samples were vortexed for 60 sec to ensure lysis of *Salmonella*. Following total RNA extraction, gDNA was removed with 4 U TURBO DNase (ThermoFisher #AM2238) per μg of RNA and then purified with RNeasy MinElute cleanup kit (Qiagen #74204). In the gDNA-free RNA, the relative human to bacterial mRNA ratio was determined by RT-qPCR measurement of human *ACTB* and *S*. Typhi *rpoD*. In brief, 150ng of RNA was reverse transcribed with iScript cDNA synthesis kit (Bio-Rad #1708891) and 1:5 dilution of this cDNA was analyzed using iTaq Universal SYBR Green Supermix (Bio-Rad #1725124) and primers listed in Table S3 on a QuantStudio 3 System (ThermoFisher). This ratio was used to combine uninfected THP-1 mRNA and late-log *S*. Typhi inoculum mRNA for the control samples following Westermann & Vogel’s approach (*43*).

### THP-1 and Intracellular *S*. Typhi Dual RNA-seq

RNA was isolated from three independent experiments and submitted to the Duke Sequencing and Genomic Technologies (SGT) Shared Resource for cDNA library preparation with Illumina Standard Total RNA Prep with Ribo-Zero Plus (Illumina #20037135). These rRNA-depleted libraries were sequenced on Ilumina NovaSeq 6000 S prime flow cell with 100 bp paired-end reads.

RNA-seq data were processed using the fastp toolkit (*44*) to trim low-quality bases and Illumina sequencing adapters from the 3’ end of the reads. Only reads that were ≥20nt after trimming were kept for further analysis. Reads were mapped to a custom genome reference combining the GRCh38v93 version of the human genome and transcriptome (*45*) with *Salmonella enterica* serovar Typhi strain Ty2 ASM754v1 genome and transcriptome using the STAR RNA-seq alignment tool (*46*). Reads were kept for subsequent analysis if they mapped to a single genomic location. Gene counts were compiled using the featureCounts tool (*47*). Only genes that had at least 10 reads in any given library were used in subsequent analysis. Normalization and differential expression within each species was carried out using the DESeq25 Bioconductor (*48*) package with the R statistical programming environment. The FDR was calculated to control for multiple hypothesis testing.

### *S*. Typhi Gene Set Enrichment Analysis (GSEA)

Intracellular *S*. Typhi RNA-seq results were converted into a ranked gene list by multiplying the log_2_(p) by the sign of the log_2_ fold-change (expression inside KO/WT THP-1). Fifteen *S*. Typhi gene sets related to divalent cation transport or virulence were generated, as shown in Table S1, and analyzed using GSEA v4.1 (*49, 50*).

### Mouse Infections

*S*. Typhimurium strains were cultured for 15 hrs overnight in 1mL Miller LB then subculture 1:33 into 1mL Miller LB for 2hr and 40min to reach SPI-1 inducing late-log growth phase (OD_600_ ∼1.8). Then bacteria washed with twice with PBS, quantified by OD_600_, and diluted to 1×10^4^ CFU/mL in sterile PBS. Litter and sex matched C57BL/6J mice bred from *Mcoln2*^+/-^ parents by the Duke DLAR breeding core were infected when 10 to 18 weeks old by intraperitoneal (IP) injection of 1,000 CFUs (100 μL) of *S*. Typhimurium DCK22 (14028s +p67GFP3.1. Inoculum was checked by plating for CFUs. Spleens were collected when the first mouse reached a humane end point four to five days post infection. The organs were homogenized by bead beating with ZrO beads (GlenMills #7305-000031) in a Mini-BeadBeater-24 (Biospec #112011). Appropriate dilutions were plated on LB agar with Amp (100 mg/mL) to calculate CFU. Infections were approved by Duke IACUC (protocol #A145-18-06).

### Statistical Analysis

Descriptive statistics were performed with GraphPad Prism v9 (GraphPad Software, US) or R v4.0.2 (R Core Team) using Hmisc (*51*) & dplyr (*52*) packages. All replication ratios were log_2_- transformed or z-scored by batch before analysis. The size of each study or number of replicates, along with the statistical tests performed can be found in figure legends. Unless otherwise indicated, all data sets passed normality tests indicating no significant deviation from a gaussian distribution. *In vitro* inter-experimental variability was removed prior to data visualization or statistical analysis by making experimental means equal to the grand mean by multiplying all values within each experiment by a normalization constant. These constants were calculated by dividing the mean of all experiments by mean of each specific experiment. Bar graphs represented the mean ± SEM (standard error of mean), unless otherwise noted. If an outlier was removed, it is noted in the figure legend along with the original value and the test used to exclude it.

## Acknowledgments

We thank the Duke University School of Medicine Sequencing and Genomic Technologies Shared Resource for providing services. We thank Kristin Cleveland and Duke DLAR Breeding Core personnel for breeding and maintenance of mouse lines. We thank the investigators and individuals from diverse populations genotyped as part of the 1000 Genomes and HapMap Projects, who have made their LCLs available through the Coriell Institute. We thank Samuel I. Miller and members of the Ko lab for useful discussion. All schematic images were generated using Biorender.com and figures were made with Adobe Illustrator v25.3.

## Author contributions

Conceptualization: KDG, LW, JSB, YC, DCK

Formal Analysis: KDG & LW

Investigation: KDG, LW, CEA, JSB, YC, MRG, DCK

Funding acquisition: KDG, DCK

Supervision: DCK

Resources: KDG, CEA, JSB, RP, DCK

Writing – original draft: KDG & DCK

Writing – review & editing: KDG, JSB, DCK

## Data and materials availability

All other data are available in the main text or the supplementary materials. GWAS summary statistics for Hi-HoST *S*. Typhi intracellular replication are available from the Duke Research Data Repository (https://doi.org/10.7924/r4x92bd76). RNA-seq data will be available in GEO upon publication.

## Supplementary Materials

Figs. S1 to S6

Tables S1 to S3

Data S1 and S2

