## SupplementalFiguresandTables for "Human variation impacting *MCOLN2* restricts *Salmonella* Typhi replication by magnesium deprivation"

**This PDF file includes:**

Figs. S1 to S6
Tables S1 to S3
Captions for Data S1 to S2

**Other Supplementary Materials for this manuscript include the following:**

Data S1 to S2

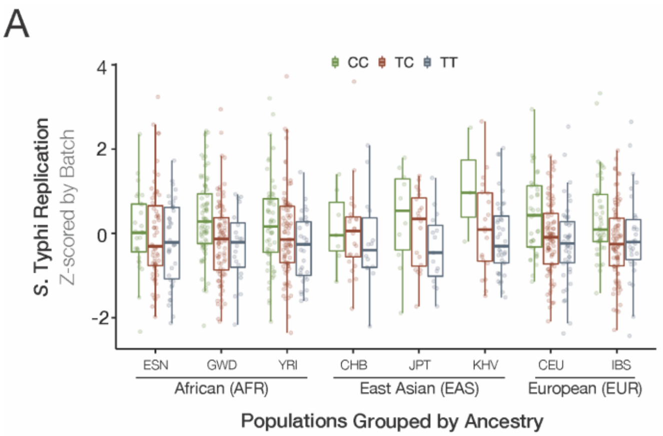


**Populations in Hi-HoST Cellular GWAS**

| Ancestry | Pop | Description | N | N |
|---|---|---|---|---|
| AFR | ESN | Esan in Nigeria | 129 | 486 |
| | GWD | Gambian Mandinka in Western Division, The Gambia | 179 | |
| | YRI | Yoruba in Ibadan, Nigeria | 178 | |
| EAS | CHB | Han Chinese in Beijing, China | 45 | 152 |
| | JPT | Japanese in Tokyo, Japan | 45 | |
| | KHV | Kihn Vietnamese in Ho Chi Minh City, Vietnam | 62 | |
| EUR | CEU | Utah residents with N & W European ancestry | 166 | 323 |
| | IBS | Iberians in Spain | 157 | |

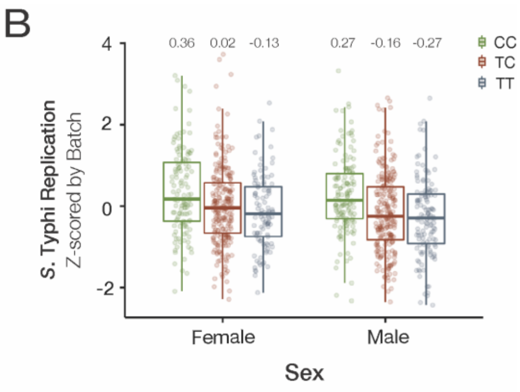


**Replciation Regressions Stratified by Sex**

| | Female | Male | All |
|---|---|---|---|
| N | 460 | 501 | 961 |
| Slope | -0.25 | -0.28 | -0.26 |
| SE of Slope | ±0.06 | ±0.06 | ±0.04 |
| p value | $1.3 \times 10^{-4}$ | $3.2 \times 10^{-6}$ | $1.7 \times 10^{9}$ |
| $R^2$ | 0.029 | 0.041 | 0.036 |

**Figure S1. The rs10873679 C-allele associates with more *S.* Typhi replication across all sampled populations and in both sexes, related to Figure 1**. (**A**) Increased z-scored *S.* Typhi replication associated with the C-allele in all populations. Individual population regressions slopes range from -0.12 in ESN to -0.40 in GWD. Table below lists number of LCLs from each population used in cellular GWAS. (**B**) The rs10873679 C-allele associated with more z-scored *S.* Typhi replication in both sexes. Linear regressions are listed in the table. A two-way ANOVA with genotype and sex finds no significant interaction between sex & *MCOLN2* genotype ($p = 0.8$) despite both individual factors being a significant source of variation (genotype $p=1.9\times10^{-9}$ & sex $p=0.02$).

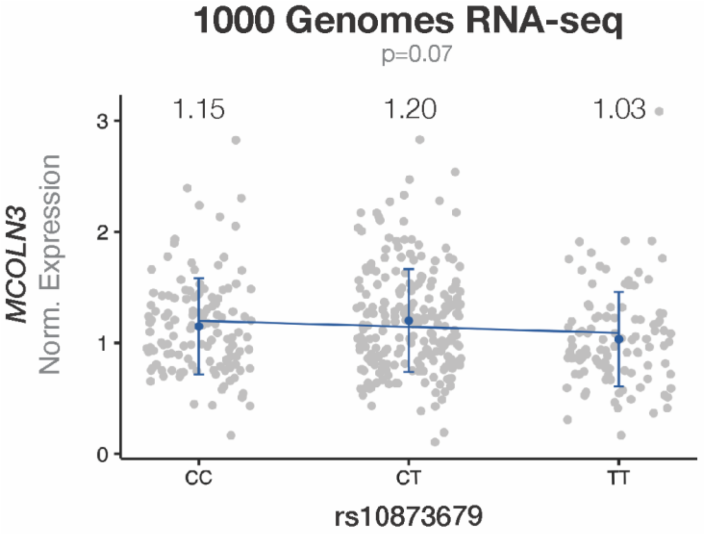


**Figure S2. rs10873679 genotype does not associate with *MCOLN3* expression in LCLs from the 1000 Genome Project**. Same dataset and analysis as Fig. 2A. The regression slope is not significantly distinguishable from zero ($\beta$ = -0.05±0.03; p = 0.07; $r^2$ = 0.005).

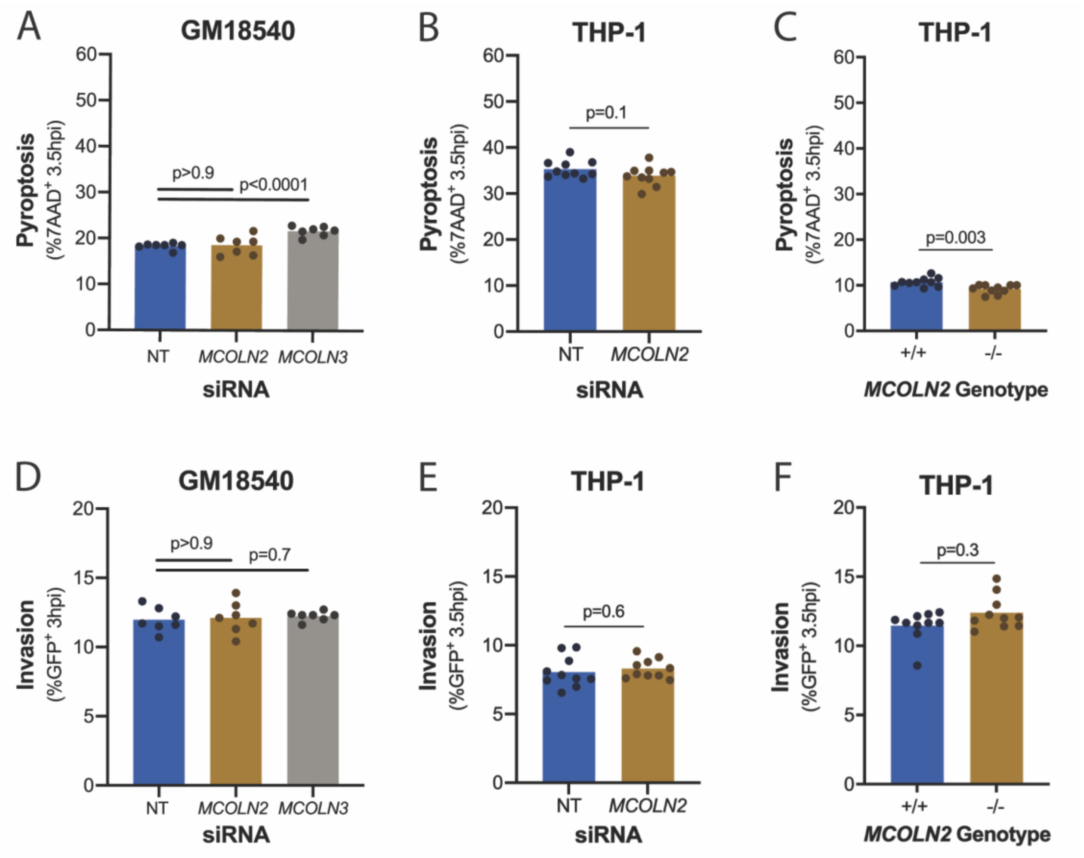


**Figure S3. Neither *MCOLN2* knockdown nor knockout affect pyroptosis or invasion during *S*. Typhi infection in Figure 2D-F.** (**A–C**) Pyroptosis is the percent of cells dead (7AAD$^{+}$) at 3.5 hpi. (**D–E**) Invasion is the percent of living (7AAD$^{-}$) and infected (GFP$^{+}$) cells at 3.5 hpi. LCL (GM18540) infection in A & D are nine replicates from three experiments from Figure 3A. THP-1 infection in B, C, E, & F are ten replicates from two experiments presented in Figure 3B & C. In A & F, the p values are from Mann-Whitney U tests. In B-E, p values are from unpaired *t* tests. While wild-type and knockout cells significantly differ in C, the effect size is only 0.5% respectively.

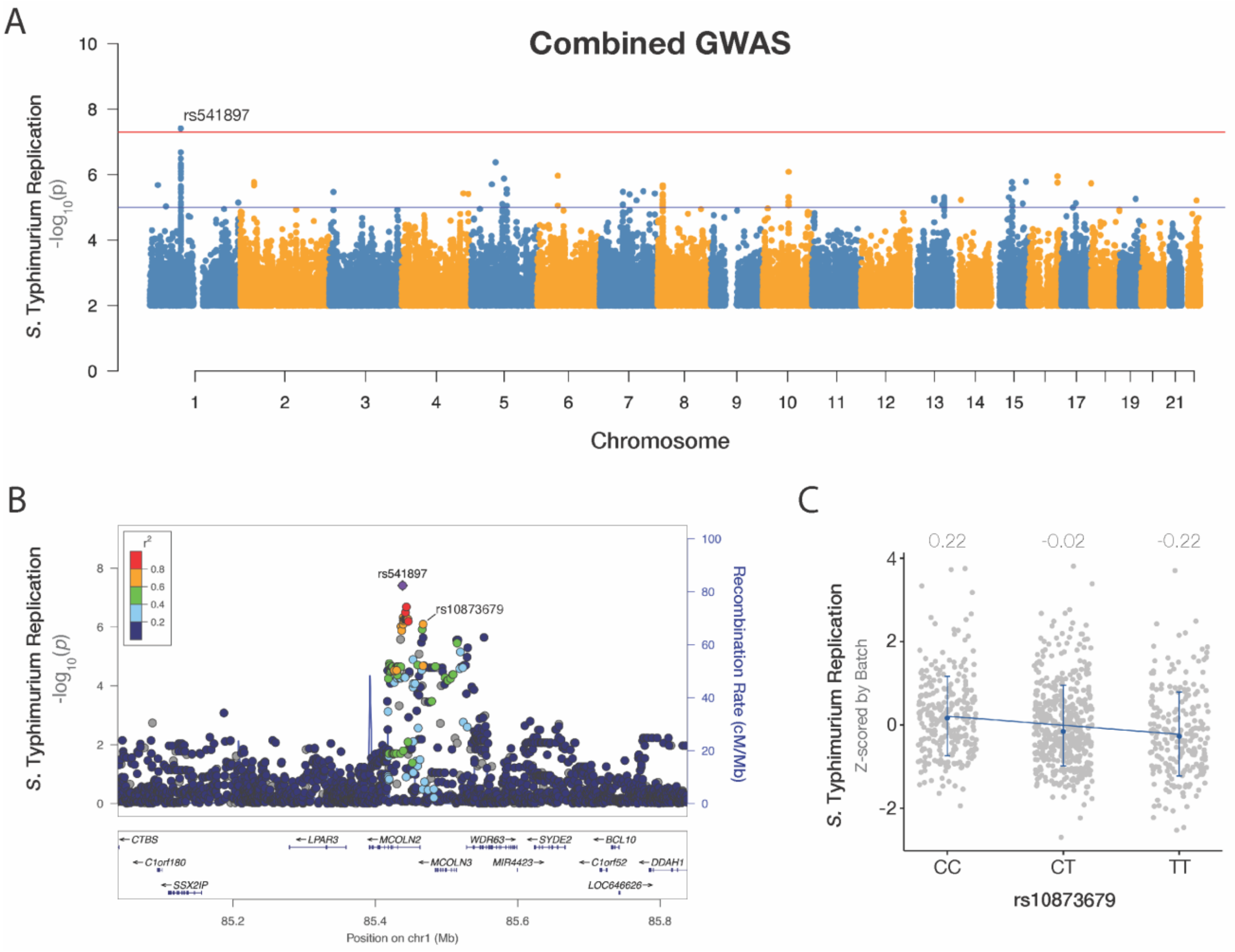


**Figure S4. Increased *S.* Typhimurium replication also associates with the rs10873679 C-allele, related to figure 2G.** (**A**) The rs541897 locus association with z-scored *S.* Typhimurium replication ($p = 3.9x10^{-8}$) passes genome-wide significance threshold ($5x10^{-8}$; red line). **(B)** The rs541897 locus is the same as the rs10873679 locus. The *S.* Typhi and *S.* Typhimurium lead SNPs are linked (LD $r^2 = 0.71$ in AFR). **(C)** The rs10873679 C-allele associates with more *S.* Typhimurium replication. The slope of a linear regression predicting replication from genotype is significantly different from zero ($\beta$=-0.22±0.04; $p = 3.5x10^{-6}$). Mean replication of each genotype is indicated above their respective dot plot. Bars are mean ±SD.

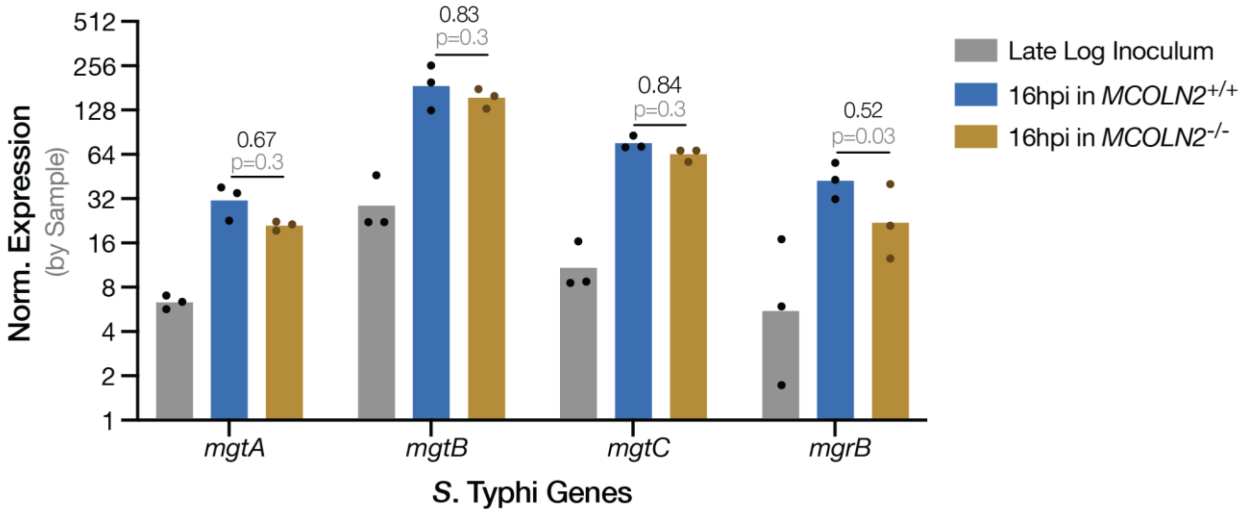


**Fig. S5.** The increased expression of $Mg^{2+}$-regulated *S*. Typhi genes inside THP-1 monocytes is consistently decreased in *MCOLN2* knockouts, same data underlying $\log_2$FC in figure 3D. Three replicates are from three experiments. Bars are geometric mean. Inoculum RNA was mixed with RNA from uninfected THP-1 cells of each genotype, so each late log replicate is the average of two technical (sequencing) replicates. Ratios (black text) are geometric mean from expression of bacteria replicating inside *MCOLN2* knockout over bacteria replicating in wild-type THP-1 cells. P values (grey) are from paired t tests corrected for multiple testing by Holm-Šídák method.

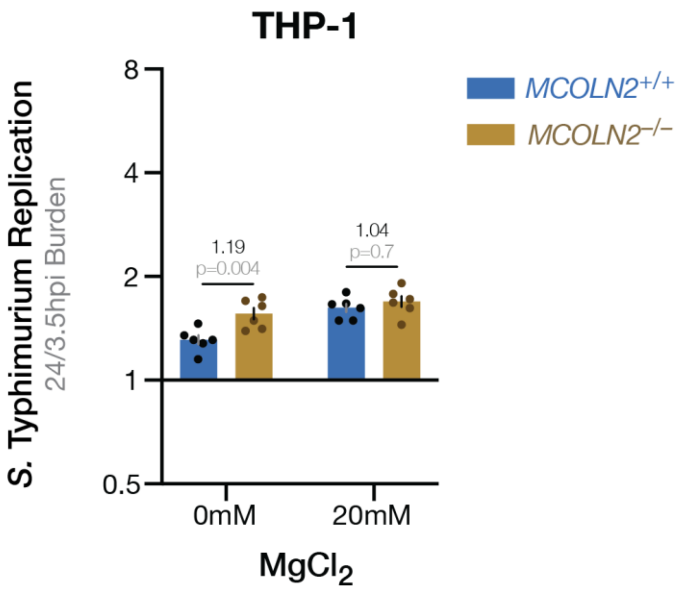


**Fig. S6.** Magnesium repletion overcomes the slight restriction of *S*. Typhimurium by MCOLN2 in THP-1 monocytes, related to figure 4A. Six replicates from two experiments. P values are from Šídák's multiple comparison following two-way ANOVA with significant main effects: *MCOLN2* genotype p=0.006, $Mg^{2+}$ repletion p=0.004, & interaction p=0.07. Like figure 4A, filter sterilized $dH_2O$ or $MgCl_2$ was added 2 hpi with MOI 10. Bars are mean ±SEM.

**Table S1.** *Salmonella* Typhi gene sets used in GSEA.

| Gene Sets | N | Genes |
|---|---|---|
| **Housekeeping** | **9** | *rpoD aroC dnaN hemD hisD sucA thrA dnaK rpoA* |
| **SPI-1** | **24** | *sicP sptP iacP prgH invA sigE invF spaM prgI invF sipA sipB sipC sipD spaS spaR spaP spaO hilA hilD spaK spaJ spaI spaH* |
| **SPI-2** | **23** | *spiC spiA ssaD ssaE sseA sseB sscA sseC sseD sseE sscB sseF sseG ssaG ssaI ssaJ ssaK ssaL ssaR ssaR ssaS ssaT ssaU* |
| **PhoP Regulon** | **19** | *pmrD mgtA phoN phoP phoQ mgtB virK ybjX mgrB pagP pagD iraP pagC mgtC pipD pagK ugtL pgtE spiC* |
| **Magnesium Influx** | **4** | *mgtA mgtB mgtS mgrB* |
| **PmrA Regulon** | **11** | *pmrC pmrA pmrB naxD arnB arnC arnA arnD arnT arnE arnF* |
| **Zinc Efflux** | **4** | *fieF zntA zntB zitB* |
| **Zinc Influx** | **4** | *zupT znuA znuB znuC* |
| **Ferrous Iron Efflux** | **2** | *fieF iceT* |
| **Ferrous Iron Influx** | **3** | *feoA feoB feoC* |
| **Manganese Influx** | **7** | *mntH sitA sitB sitC sitD mntR mntS* |
| **Copper Response** | **8** | *copA cueR cueO cueP scsA scsB scsC scsD* |
| **MFS Efflux** | **5** | *mdtH mdtM mdtG emrD mdfA* |
| **TolC Efflux** | **5** | *macB tolC acrA acrE marA* |
| **RpoE Regulon** | **5** | *rpoE rseA rseB rseC* |

**Table S2.** *Salmonella enterica* Typhi strains used in this study.

| Serovar | Designation | Genotype | Plasmid | Resistance | Derived From |
|---|---|---|---|---|---|
| *S.* Typhi | DCK33 | Ty2 | p67GFP3.1 | Amp | CS092 |
| | CS021 | Ty2 Δ*phoPQ* (956bp deletion) | p67GFP3.1 | Amp | Ty800 |
| | DCK723 | Ty2 Δ*ssaT* | p67GFP3.1 | Amp | DCK722 |
| | DCK1080 | Ty2 Δ*mgtB* | – | – | DCK165 |
| | DCK1120 | Ty2 Δ*mgtA* Δ*mgtB* | – | – | DCK1080 |
| | DCK1122 | Ty2 Δ*mgtA* Δ*mgtB* | p67GFP3.1 | Amp | DCK1120 |
| *S.* Typhimurium | DCK22 | 14028s | p67GFP3.1 | Amp | CS093 |
| | DCK483 | 14028s | pWSK29 | Amp | CS093 |
| | DCK484 | 14028s | pWSK129 | Kan | CS093 |

**Table S3.** Oligos grouped by function in study.

***Lambda-Red Recombination***

| Target Gene | Gene ID | Primer Designations | Fwd for Kan$^R$ Cassette Generation | Rvr for Kan$^R$ Cassette Generation |
|---|---|---|---|---|
| *mgtA* | t4491 | DK972 & DK973 | TATAATCCGCGGCGCAAATTATTTACTTACCGGAGGCGAC gtgtaggctggagctgcttc | TCGGGGATTAAGCACGCTGGCGAATCCCCGACGAAAGTGT catatgaatatcctccttag |
| *mgtB* | t3755 | DK974 & DK975 | ATATGCAGGAAACACTACACCTTAATTTTGGGGATTCATC gtgtaggctggagctgcttc | TATCGGGTGAGCGATTCATCTGGGCGATCCTCAAACATTA catatgaatatcctccttag |
| *ssaT* | t1289 | DK3 & DK433 | tacccggcagataatgttacgaattggagagcatggttga gtgtaggctggagctgcttc | TCACGTAATTTCTTTTCTGTAGGCTGTTCTGTTTTCTCGC catatgaatatcctccttag |

***Deletion Confirmation***

| Target Gene(s) | Gene ID | Primer Designations | Fwd | Rvr |
|---|---|---|---|---|
| *phoPQ* | t1689 & t1690 | DK1008 & DK1009 | CATGACGCCGGCAAATTATATC | GAAAGTCGGGCCAGTTAAGA |
| *ssaT* | t1289 | DK434 & DK435 | CGGTAGTTGGTGTCATCGTAAG | AGCGCAATCAGCTGAAATAATG |
| *mgtA* | t4491 | DK980 & DK981 | CGTGACGCTGATGGTGATAAA | ACATCTCCTCCTCTCGTTCTG |
| *mgtB* | t3755 | DK982 & DK983 | CAGGCGTATAAGGAGGGAATG | ACACCAACAGGCTAATCAGTAA |
| *MCOLN2* | ENSG00000153898 | DK859 & DK860 | CTGGGGTAATTTTCCAAAAGCAGT | AGGTTATTCTTCTGTGGGACTTGT |

***SYBR RT-qPCR***

| Target Gene(s) | Gene ID | Primer Designations | Fwd | Rvr |
|---|---|---|---|---|
| *ACTB* | ENSG00000075624 | DK880 & DK881 | CCTGTACGCCAACACAGTGC | ATACTCCTGCTTGCTGATCC |
| *rpoD* | t3131 | DK628 & DK629 | GAAATGGGCACTGTTGAACTG | CAGATAGGTAATGGCTTCCGG |

**Data S1.** (separate file) Intracellular replication measurements from 961 LCLs.

**Data S2.** (separate file) Transcriptomic data of *S.* Typhi within wild-type and *MCOLN2-/-* THP-1 monocytes.
